# RABL6A promotes pancreatic neuroendocrine tumor angiogenesis and progression *in vivo*

**DOI:** 10.1101/2021.03.17.435790

**Authors:** Chandra K. Maharjan, Courtney A. Kaemmer, Viviane P. Muniz, Casey Bauchle, Sarah L. Mott, K.D. Zamba, Patrick Breheny, Mariah R. Leidinger, Benjamin W. Darbro, Samuel B. Stephens, David K. Meyerholz, Dawn E. Quelle

**Affiliations:** Department of Neuroscience and Pharmacology, The University of Iowa, Iowa City, Iowa; Fraternal Order of Eagles Diabetes Research Center, The University of Iowa, Iowa City, Iowa; Holden Comprehensive Cancer Center, The University of Iowa, Iowa City, Iowa; Department of Biostatistics, The University of Iowa, Iowa City, Iowa; Department of Pathology, The University of Iowa, Iowa City, Iowa; Department of Pediatrics, Carver College of Medicine, The University of Iowa, Iowa City, Iowa

**Keywords:** Pancreatic neuroendocrine tumors, RIP-Tag2 mouse model, RABL6A, angiogenesis, c-Myc

## Abstract

Pancreatic neuroendocrine tumors (pNETs) are difficult-to-treat neoplasms whose incidence is rising. Greater understanding of pNET pathogenesis is needed to identify new biomarkers and targets for improved therapy. RABL6A, a novel oncogenic GTPase, is highly expressed in patient pNETs and required for pNET cell proliferation and survival *in vitro*. Here, we investigated the role of RABL6A in pNET progression *in vivo* using a well-established model of the disease. RIP-Tag2 (RT2) mice develop functional pNETs (insulinomas) due to SV40 large T-antigen expression in pancreatic islet β cells. RABL6A loss in RT2 mice significantly delayed pancreatic tumor formation, reduced tumor angiogenesis and mitoses, and extended survival. Those effects correlated with upregulation of anti-angiogenic p19ARF and downregulation of proangiogenic *c-Myc* in RABL6A-deficient islets and tumors. Our findings demonstrate that RABL6A is a bona fide oncogenic driver of pNET angiogenesis and development *in vivo*.

## Introduction

Pancreatic neuroendocrine tumors (pNETs) are incurable, typically indolent neoplasms originating from multiple neuroendocrine cell types, particularly β cells, within the islets of Langerhans (Scott & Howe, 2019). In healthy individuals, pancreatic β cells maintain proper blood glucose levels via insulin production and release. β cell dysfunction leads to diabetes while their neoplastic transformation results in pNET formation. Although pNETs are uncommon, their incidence has risen more than 4-fold over the past four decades (Dasari et al., 2017; Halfdanarson, Rabe, Rubin, & Petersen, 2008; Sun, 2017). This is alarming because the molecular etiology of pNET development is only partly understood and current therapies have little to no impact on improving overall patient survival. Greater insight into essential mechanisms driving this disease will pave the way for more effective targeted therapies.

We recently identified a new potential driver of pNETs, a RAB-like GTPase named RABL6A. The protein was first discovered as a novel binding partner of the Alternative Reading Frame (ARF; p14ARF in humans, p19ARF in mice) tumor suppressor (Tompkins, Hagen, Zediak, & Quelle, 2006). Later studies showed RABL6A is essential for the growth and survival of various tumor types, controls central cancer pathways (RB1, p53, PP2A, AKT, mTOR, ERK), and is a biomarker of poor survival in several cancers including breast and pancreatic adenocarcinomas (Feng et al., 2020; Hagen et al., 2014; Kohlmeyer et al., 2020; Li et al., 2013; Lui et al., 2013; Montalbano, Lui, Sheikh, & Huang, 2009; Muniz et al., 2013; H. Tang et al., 2016; Umesalma et al., 2019). In pNETs, RABL6A is highly expressed at the genetic and protein levels in human primary and metastatic tumors (Hagen et al., 2014; Scott et al., 2020). Functional studies in cultured pNETs revealed RABL6A is required for pNET cell proliferation and viability (Hagen et al., 2014; Umesalma et al., 2019). Moreover, RABL6A acts through several clinically relevant pNET pathways. This includes inhibition of the retinoblastoma (RB1) and PP2A tumor suppressors and activation of oncogenic AKT and mTOR (Hagen et al., 2014; Umesalma et al., 2019).

In this study, we explored the *in vivo* oncogenic role of RABL6A in pNET development and progression. We employed the RIP-Tag2 (RT2) transgenic mouse model, which has been used extensively to examine pNET biology and evaluate the utility of promising new therapies (Bergers, Song, Meyer-Morse, Bergsland, & Hanahan, 2003; Casanovas, Hicklin, Bergers, & Hanahan, 2005; Chiu, Nozawa, & Hanahan, 2010; Christofori, Naik, & Hanahan, 1994; Hanahan, 1985; Hanahan & Folkman, 1996; Inoue, Hager, Ferrara, Gerber, & Hanahan, 2002; Olson, Chu, Perry, Nolan-Stevaux, & Hanahan, 2011; Parangi, Dietrich, Christofori, Lander, & Hanahan, 1995; Sodir et al., 2011; Ulanet & Hanahan, 2010). In RT2 mice, the rat insulin promoter (RIP) drives expression of the SV-40 large T-antigen in pancreatic β cells, causing their oncogenic transformation in a highly reliable, sequential, and age-dependent manner (Hanahan, 1985). We found that loss of RABL6A in this pNET model significantly reduced the tumor phenotype and increased survival, demonstrating a critical *in vivo* role for RABL6A in pNET pathogenesis.

## Results

### Description and generation of the mouse models

RT2 mice exhibit temporally defined, multi-stage development of pNETs through oncogenic transformation of the islet β cells (Hanahan, 1985). Over half of the normal pancreatic islets in RT2 mice proliferate aberrantly and transform into hyperplastic islets by 3-5 weeks. Of the hyperplastic islets, about one-fifth undergo an angiogenic switch and become vascularized angiogenic islets by 7-9 weeks, with one-fourth of those progressing into enlarged pNETs starting at 10-12 weeks (Christofori et al., 1994; Folkman, Watson, Ingber, & Hanahan, 1989; Hanahan, 1985). The pNETs that arise are functional insulinomas that secrete insulin.Representative hyperplastic islets, red angiogenic islets, and red vascularized pNETs isolated by pancreatic perfusion from 12-week-old RT2 male mice are shown in ***Figure 1A***.

**Figure 1.**
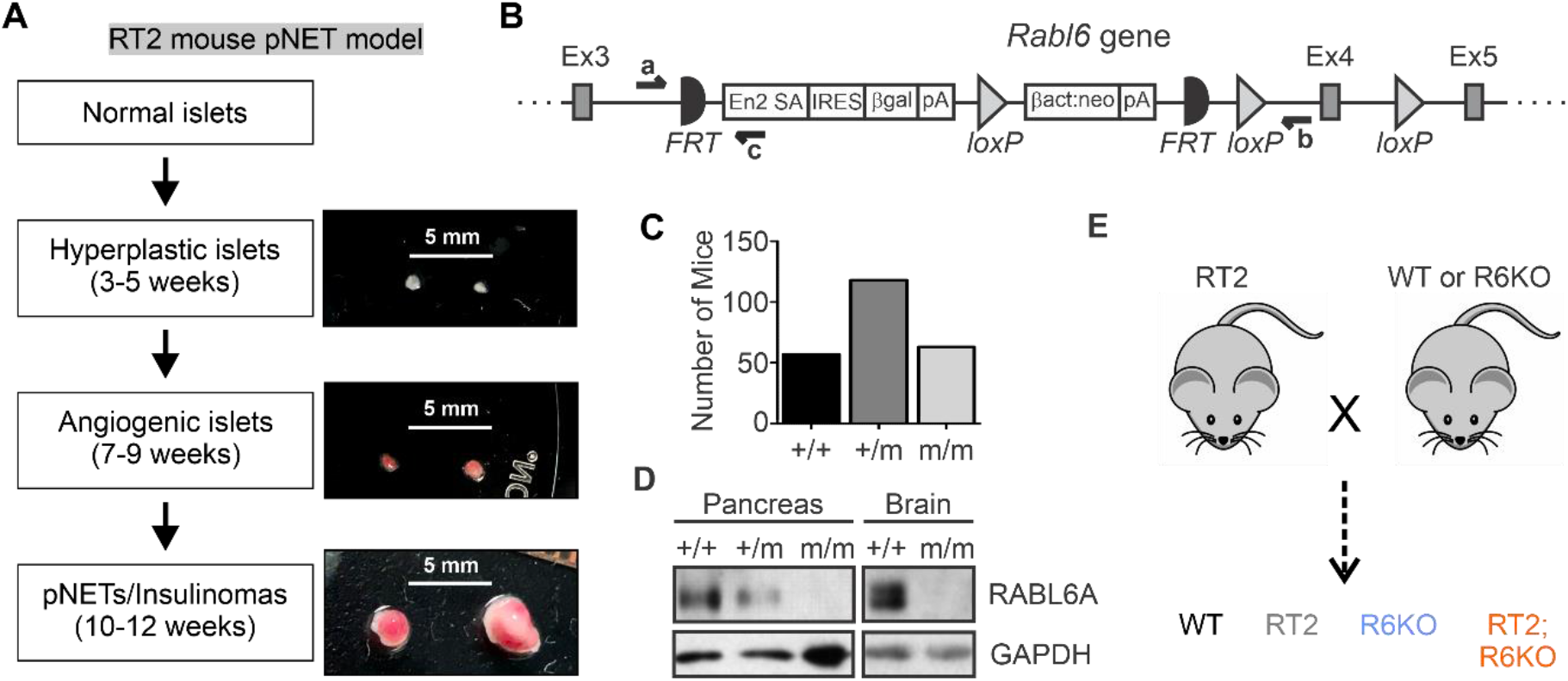
Mouse models to study the *in vivo* role of RABL6A in pNET pathogenesis (A) Schematic outlining the sequential progression of islet transformation into functional pNETs (insulinomas) in RT2 mice. Time in parentheses represent the age of RT2 mice when the indicated islets are first observed. Right; Images of each type of lesion (with scale bars) isolated from a 12-week-old RT2 male. (B) Diagram of the targeted *Rabl6* gene locus (exons 3-5 shown) in the gene-trap knock-out mouse model. Indicated sites include FRT (Flippase recombinase), loxP (Cre-recombinase), and primers for detecting endogenous *Rabl6* allele (‘a’ and ‘b’) and mutant (m) allele (‘a’ and ‘c’). (C) Number of mice with +/+, +/m, or m/m genotype from *Rabl6^+/m^* heterozygous mouse crossings showing a Mendelian distribution of 1:2:1. (D) Western blots showing loss of RABL6A protein in *Rabl6^m/m^* mouse pancreas and brain. (E) Simplified schematic of RT2 crosses with WT or *Rabl6^m/m^* (R6KO) mice over multiple generations to obtain the experimental cohorts (WT, RT2, R6KO, and RT2; R6KO). **Figure supplement 1.** Genotyping RIP-Tag2 (RT2) and R6KO mice.

To examine the oncogenic role of RABL6A in pNET development, we first generated *Rabl6* knockout (R6KO) mice. *Rabl6* chimeric mice were obtained from the Knockout Mouse Project (KOMP) and bred with C57BL/6N animals to achieve germline transmission. A gene trap sequence consisting a splice acceptor and a poly(A) adenylation site was targeted between 5’ exons 3 and 4 of the *Rabl6* gene *(**Figure 1B***), interfering with normal gene splicing and expression. After successful germline transmission, the heterozygous *Rabl6^+/m^* mutant mice were interbred to obtain homozygous *Rabl6^m/m^* mice, henceforth referred to as RABL6A knockout (R6KO). A classical 1:2:1 Mendelian ratio was obtained from intercrosses of *Rabl6^+/m^* mice demonstrating that RABL6A deficiency is not embryonically lethal *(**Figure 1C***). Western analysis verified loss of RABL6A protein expression in homozygous R6KO pancreatic and brain lysates *(**Figure 1D***).

RT2 and R6KO mice were crossed for multiple generations to obtain double transgenic RT2; R6KO mice as well as RT2, R6KO and wild-type (WT) controls *(**Figure 1E**;* genotyping results shown in ***Figure 1- figure supplement 1***).

### RABL6A expression reduces survival in RT2 mice

RT2 mice have a short life expectancy due to excessive insulin secretion from the pNETs that induces hypoglycemic shock and death (Hanahan, 1985). We monitored the survival of male and female mice within all four experimental cohorts to determine if RABL6A loss improves survival in RT2 mice. All WT and R6KO mice of both sexes were alive throughout the entire observation period. A significant difference was seen in the survival of RT2 females (median 13.3 weeks) versus RT2 males (median 15.7 weeks) (***Figure 2***), an observation that we could not find noted in prior reports. As hypothesized, loss of RABL6A significantly increased the survival of RT2; R6KO males (median 18.0 weeks). A modest increase in the median survival of RT2; R6KO females (14.1 weeks) was seen relative to that of RT2 females (13.3 weeks), but the difference was not statistically significant (***Figure 2; Table 1***). After adjusting for sex, an overall difference in survival was evidenced between RT2 and RT2; R6KO mice with RT2 mice being at 59% increased risk of death (HR=1.59, p<0.05).

**Figure 2.**
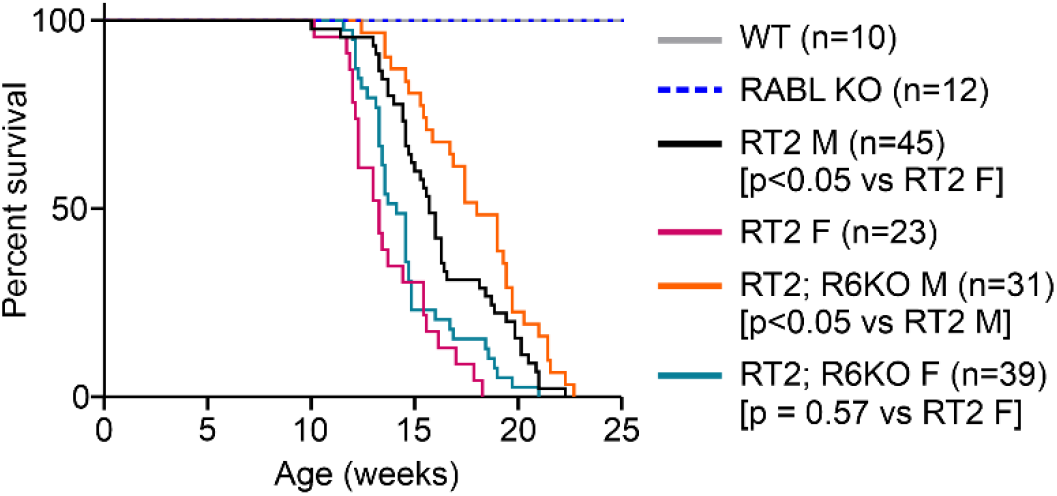
Loss of RABL6A improves survival of RT2 mice. Kaplan-Meier survival curves of overall survival for the indicated mice through 25 weeks after birth. All WT and R6KO mice were healthy and alive during this period. The Kaplan-Meier method was used to estimate the survival curves and group comparisons were made using the log rank test. P values for the indicated comparisons are shown; (n), number of animals in each cohort. **Source data 1**. Fig 2 source data.

**Table 1.** Summary of biological effects caused by RABL6A loss in RT2 mice.

| Analyses |  | Females |  |  | Males |  |  |
| --- | --- | --- | --- | --- | --- | --- | --- |
|  |  | RT2 | RT2; R6KO | p-value | RT2 | RT2; R6KO | p-value |
| Survival (wks) | Median survival | 13.3 | 14.1 | 0.57 | 15.7 | 18.0 | <0.05 |
| pNET formation | Tumor burden (mm<sup>3</sup>) | 8.2±2.7 | 0.7±0.5 | <0.05 | 52.0±14.2 | 16.1±3.8 | <0.05 |
|  | % Endocrine area | 9.3±2.4 | 2.5±0.6 | <0.05 | 17.7±2.5 | 13.5±1.9 | 0.11 |
| Islet transformation | % Angiogenic islets | 36.7±3.3 | 14.2±2.5 | <0.05 | 31.4±4.5 | 21.9±5.0 | 0.19 |
|  | % Vasculature area | 8.9±1.1 | 4.2±1.0 | <0.05 | 7.4±1.0 | 7.3±1.3 | 0.95 |
|  | Mitoses (per field of view at 600x) | 1.9±0.2 | 0.9±0.2 | <0.05 | 3.0±0.3 | 2.4±0.2 | 0.32 |
| c-Myc expression | c-Myc mRNA | 1.1±0.2 | 0.6±0.1 | <0.05 | 1.7±0.6 | 0.5±0.1 | 0.11 |
|  | c-Myc protein | ND | ND | ND | 1.0±0.6 | 0.7±0.2 | 0.28 |
| ARF expression | p19ARF protein | ND | ND | ND | 1.0±0.3 | 4.2±1.0 | <0.05 |

Data represent mean ± SEM obtained from 10-week-old females and 12-week-old males (except for the survival analysis). p-values indicative of statistically significant differences are shown in blue. ND, not determined.

### RABL6A increases tumor burden in RT2 mice in an age-dependent manner

Given the sex-dependent differences in overall survival for RT2 and RT2; R6KO mice, we evaluated the development and progression of tumors according to different timelines for male and female animals. Female cohorts were euthanized for histopathologic and molecular analysis at 8, 10, and 12 weeks of age whereas males were euthanized at 8, 12, and 16 weeks of age (***Figure 3A***). At each time point, pancreatic islets and tumors were isolated via pancreatic perfusion or the whole pancreas was excised for fixation and histopathology.

**Figure 3.**
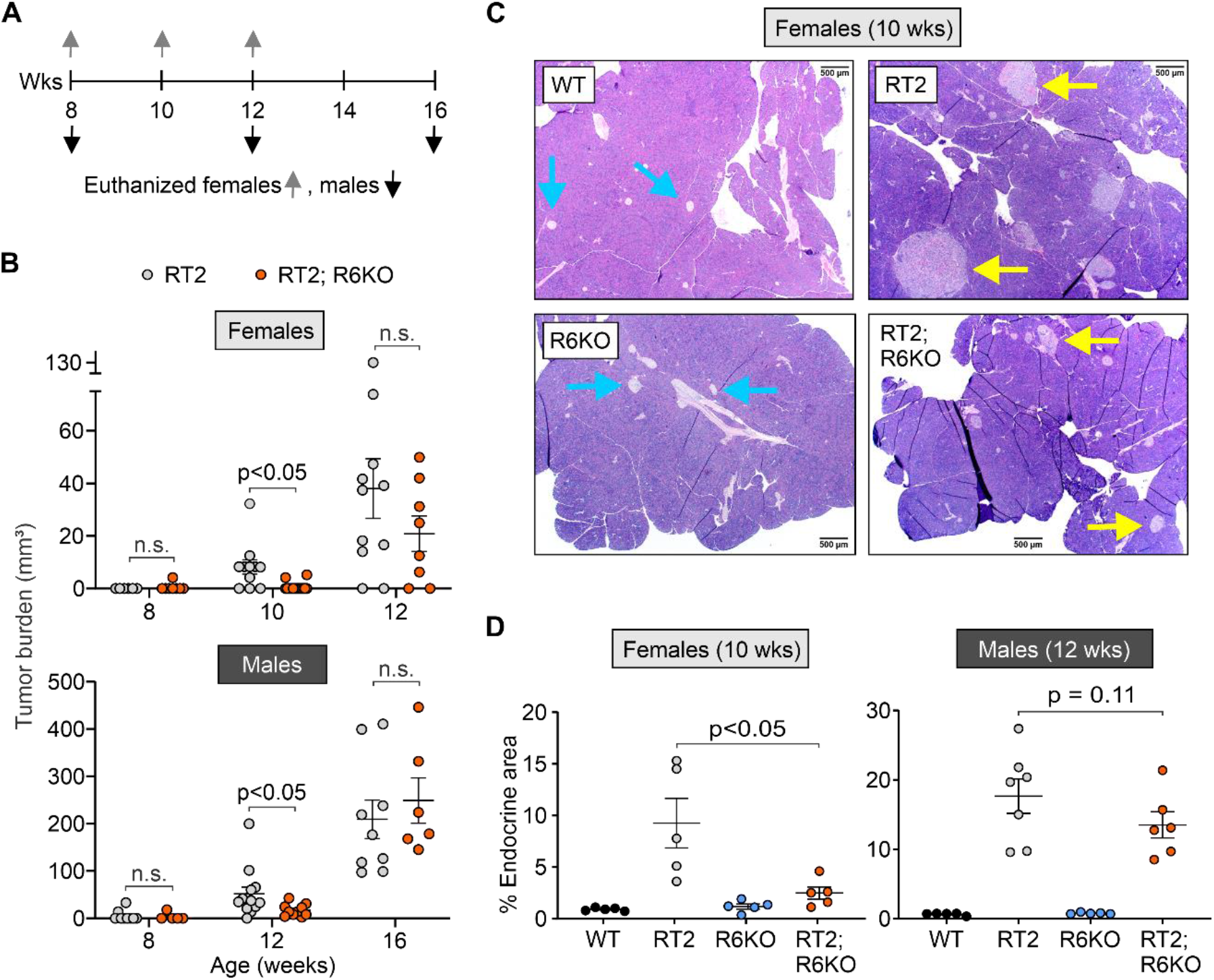
Loss of RABL6A delays tumor formation in RT2 mice. **(A)** Study timeline showing ages in weeks (Wks) of female (up arrow) and male (down arrow) mice when they were euthanized for islet isolation or histopathology of the pancreas. **(B)** Tumor burden is significantly reduced in 10-week-old female and 12-week-old male RT2; R6KO mice (orange) compared to age-matched RT2 controls (gray). P values shown; n.s., not significant. N = Females: 8 weeks (RT2 = 6, RT2; R6KO = 8), 10 weeks (RT2 = 11, RT2; R6KO = 14), 12 weeks (RT2 = 11, RT2; R6KO = 8); Males: 8 weeks (RT2 = 9, RT2; R6KO = 5), 12 weeks (RT2 = 13, RT2; R6KO = 11), 16 weeks (RT2 = 9, RT2; R6KO = 6). **(C)** Representative images of H&E-stained pancreas from 10-week-old WT, RT2, R6KO, and RT2; R6KO females. Blue arrows, normal islets; yellow arrows, transformed islets. Scale bar = 500 μM. **(D)** Reduction in the % endocrine area in 10-week-old RT2: R6KO females (left, p<0.05) with similar trend (p=0.11) in 12-week-old RT2; R6KO males (right) versus RT2 controls. Data in A, B and D represent the mean +/- SEM with each individual animal represented by a dot. N = Females (WT = 5, RT2 = 5, R6KO = 5, RT2; R6KO = 5); Males (WT = 5, RT2 = 7, R6KO = 5, RT2; R6KO = 6). Linear regression models were used to evaluate differences in tumor burden and beta regression models to compare percent endocrine area. **Source data 1:** Figure 3B source data. **Source data 2:** Figure 3D source data. **Figure supplement 1.** Plasma insulin concentration over time. **Figure supplement 2.** Percent endocrine area in females (8 and 12 weeks) and males (8 and 16 weeks).

We first quantified the volume of tumors harvested from RT2 and RT2; R6KO mice and observed an age-dependent decrease in tumor burden in mice lacking RABL6A (***Figure 3B; Table 1***). Specifically, 10-week-old female and 12-week-old male RT2; R6KO mice had significantly lower tumor burden compared to their RT2 counterparts. The difference in tumor burden associated with RABL6A loss was absent in older animals, reflecting a delay rather than abolishment of tumor formation (***Figure 3B***). Since pNETs in RT2 mice cause hyperinsulinemia, we tested whether plasma insulin levels correlated with tumor burden (***Figure 3- figure supplement 1***). WT and R6KO mice, which both lack tumors, exhibited normal, low plasma insulin concentrations (<2 ng/mL) throughout the observation timeframe. Notably, R6KO male mice had reduced insulin levels compared to WT mice as the animals aged. By comparison, plasma insulin levels increased at similar rates in both RT2 and RT2; R6KO mice (males and females) over time regardless of differences in tumor burden.

Complementary histopathological analyses of H&E stained pancreata from each experimental cohort enabled further assessment of tumor burden. WT and R6KO mice had small, normal looking islets (***Figure 3C***) that represented a small fraction of the entire pancreas, as measured by the percent endocrine area (cumulative area of islets relative to the total pancreatic area) (***Figure 3D***). The pancreas of RT2 males and females contained enlarged islets (comprising hyperplastic islets, angiogenic islets, and tumors), which were invasive on the surrounding exocrine tissue (***Figure 3C***). Quantitative analysis revealed remarkably high percentage endocrine area in RT2 mice compared to WT and R6KO animals. The 8- and 10-week-old RT2; R6KO females had smaller islets and significantly reduced percent endocrine area compared to age-matched RT2 counterparts (***Figure 3C, D; Table 1; Figure 3- figure supplement 2***). A lower average percent endocrine area was likewise observed in 12-week-old RT2; R6KO males (13.5%) compared to RT2 males (17.7%), albeit smaller in magnitude and not statistically significant (***Figure 3D; Table 1; Figure 3- figure supplement 2***).

### RABL6A promotes angiogenesis during early stages of pNET formation

In RT2 mice, the progression of hyperplastic islets to angiogenic islets, referred to as angiogenic switch, is a key event in pNET development (Hanahan & Folkman, 1996). p19^ARF^, a well-known tumor suppressor and binding partner of RABL6A, was previously shown to disrupt the angiogenic switch and tumor initiation in RT2 mice (Ulanet & Hanahan, 2010). To determine if RABL6A promotes pNET angiogenesis in RT2 mice, we compared the percentage of angiogenic islets and their vasculature in H&E-stained RT2 and RT2; R6KO pancreatic sections (***Figure 4***).

**Figure 4.**
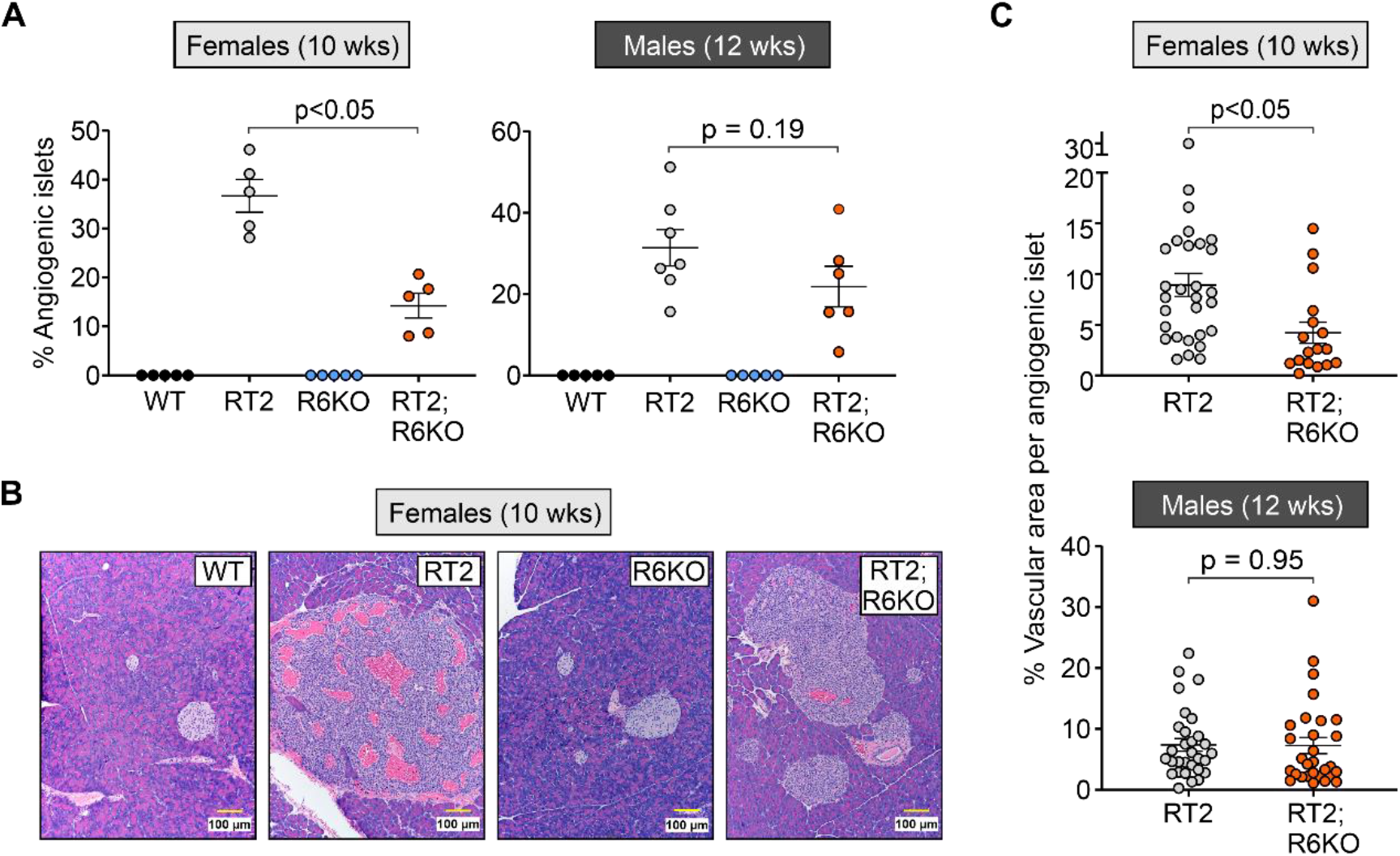
Loss of RABL6A reduces the angiogenic switch in RT2 mice. **(A)** Decreased percent angiogenic islets in 10-week-old female RT2; R6KO mice relative to RT2 controls. 12-week-old males trended similarly without statistical significance. Each dot equals data for an individual mouse pancreas. **(B)** Representative images of normal islets in 10-week-old female WT and R6KO mice versus angiogenic islets or tumors in RT2 and RT2; R6KO mice. Scare bar = 100 μm. **(C)** Reduction in percent vascular area per angiogenic islet or tumor in RT2; R6KO 10-week-old females. WT and R6KO islets lack vasculature and are not shown. Each dot represents an individual angiogenic islet or tumor. Data in A and C are reported as the mean +/- SEM. N = Females (WT = 5, RT2 = 5, R6KO = 5, RT2; R6KO = 5); Males (WT = 5, RT2 = 7, R6KO = 5, RT2; R6KO = 6). Beta regression model was used to evaluate differences in percent angiogenic islets and vascular area per angiogenic islet. Random intercepts were included to account for the correlated nature of angiogenic islets within the same mouse in the evaluation of the vascular area per angiogenic islet. **Source data 1:** Figure 4A source data. **Source data 2:** Figure 4C source data. **Figure supplement 1.** Percent angiogenic islets and vascular area per angiogenic islet in females (8 and 12 weeks) and males (8 and 16 weeks).

Angiogenic islets, characterized by pockets of blood vessels (capillaries not counted) contained within their circumference, were absent in control WT and R6KO pancreas (***Figure 4A, B***). In RT2 mice, the number of angiogenic islets relative to the total islets increased with age (***Figure 4A, Figure 4- figure supplement 1***). In female mice, 10-week-old RT2 females displayed a significantly higher percentage of angiogenic islets compared to their RT2; R6KO counterparts (***Figure 4A, B; Table 1; Figure 4- figure supplement 1***). While the differences between 12-week-old male cohorts did not reach statistical significance, the average percentage of angiogenic islets was similarly greater in RT2 mice relative to RT2; R6KO animals (31.4% to 21.9%) (***Figure 4A; Table 1***).

We then asked whether RABL6A deficiency attenuates the extent of vasculature within angiogenic islets in RT2 mice. For this, we quantified percentage area of vasculature in 3-6 representative angiogenic islets in each RT2 or RT2; R6KO pancreas. RT2 angiogenic islets were highly vascularized and the extent of vasculature was significantly reduced in the angiogenic islets of 10-week-old RT2; RABLKO females (***Figure 4B, C; Table 1***). This effect was not observed in RT2; R6KO males compared to age matched RT2 males at 8, 12 or 16 weeks (***Figure 4C; Table 1; Figure 4- figure supplement 1***). These cumulative findings demonstrate that RABL6A promotes the angiogenic switch and increases vascularization during the early stages of pNET pathogenesis in RT2 females with a less pronounced effect (without statistical significance) in RT2 males.

### RABL6A drives pNET cell mitosis early in tumor development

RABL6A promotes pNET cell proliferation *in vitro* (Hagen et al., 2014; Umesalma et al., 2019). Mitotic counts on H&E sections are one of the grading parameters for pNETs (Rindi et al., 2018). Thus, we scored the number of mitoses (cells with condensed nuclear DNA) in nonoverlapping high-power fields of pancreatic islets and tumors in RT2 and RT2; R6KO animals (***Figure 5; Figure 5- figure supplement 1***). A significant reduction in mitoses was seen in both 8- and 10-week-old female RT2; R6KO islets compared to those in age-matched RT2 mice (***Figure 5A, B; Table 1; Figure 5- figure supplement 1***). No significant differences in mitotic counts were seen in the islets and pNETs of older (12-week) female RT2; R6KO and RT2 mice or in 8-week-old male cohorts (***Figure 5- figure supplement 1***). However, 12-week-old RT2; R6KO males displayed a modest decrease in mitoses (mean 2.4 vs 3.0 in RT2 controls; not significant) that mirrored the differences in younger female cohorts (***Figure 5A, B; Table 1; Figure 5- figure supplement 1***). Of note, 16-week-old RT2; R6KO males showed elevated pNET mitoses relative to age-matched RT2 controls (***Figure 5- figure supplement 1***), possibly reflecting compensatory hyperactivation of proliferative signaling following sustained loss of RABL6A (Umesalma et al., 2019). Together, these results suggest RABL6A enhances islet cell mitosis at early time points in PNET development, with effects most evident in female mice.

**Figure 5.**
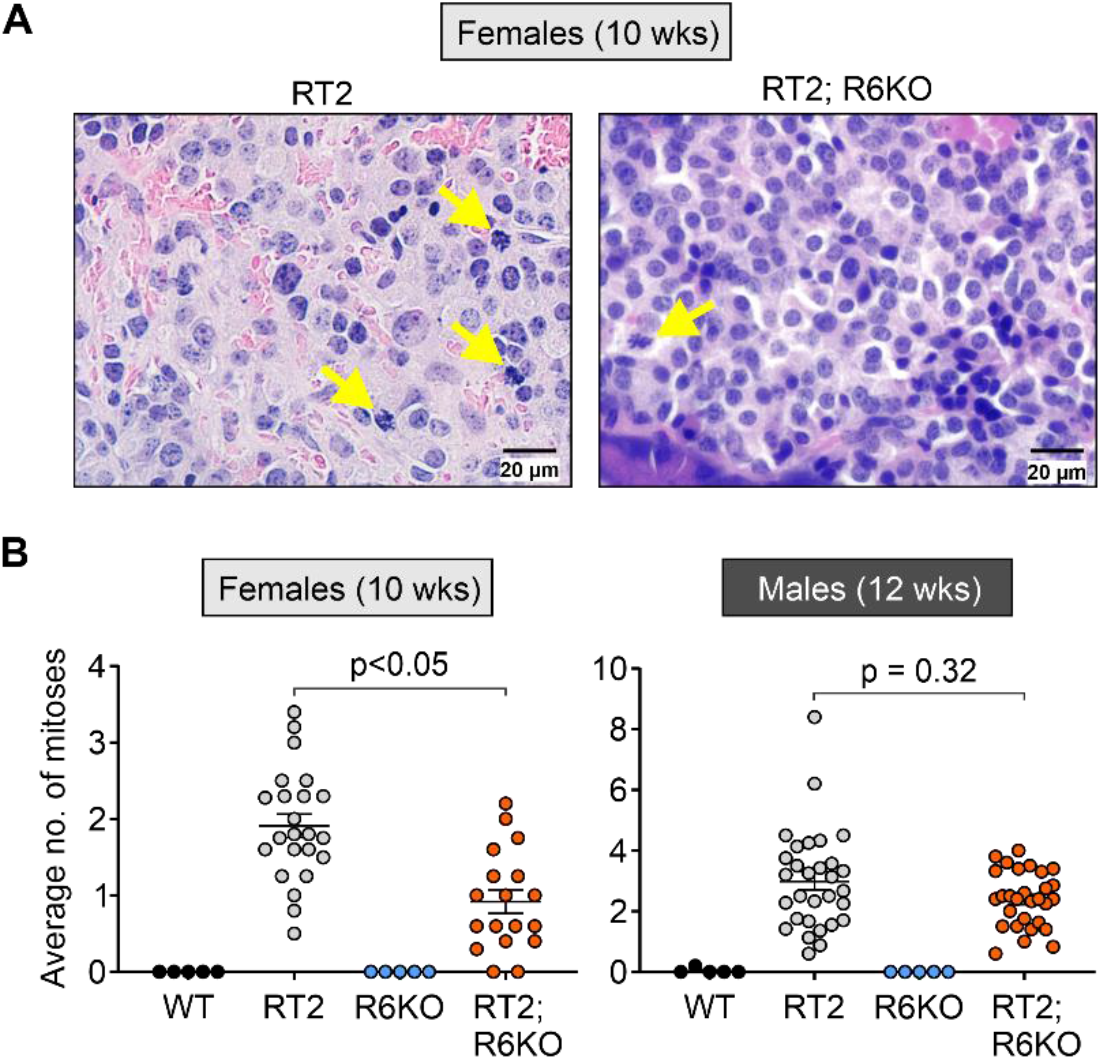
Loss of RABL6A reduces islet cell mitoses in RT2 female mice. Mitoses were counted on 3-6 random fields of view within an islet at 600x magnification and their average was calculated. **(A)** Representative images of RT2 and RT2; R6KO islet sections. Arrows indicate mitoses. Scale bar = 20 μM.**(B)** The average number of mitoses (per field) is significantly reduced in the transformed islets of 10-week-old RT2; R6KO females (left) compared to age-matched RT2 females. Data are the mean +/- SEM with each dot representing an average number of mitoses per field of vision for an individual islet. N = Females (WT = 5, RT2 = 5, R6KO = 5, RT2; R6KO = 5); Males (WT = 5, RT2 = 7, R6KO = 5, RT2; R6KO = 6). Linear regression was used to compare average number of mitoses between groups. **Source data 1:** Figure 5B source data. **Figure supplement 1.** Average number of mitoses in females (8 and 12 weeks) and males (8 and 16 weeks).

### RABL6A promotes *c-Myc* and limits ARF expression in mouse pancreatic islets

Targeted overexpression of ectopic c-Myc in pancreatic β cells yields highly angiogenic islet tumors in mice while its deactivation induces tumor regression preceded by vascular degeneration and β cell apoptosis (Pelengaris, Khan, & Evan, 2002). Other work in RT2 mice showed that endogenous c-Myc activity is required for maintaining pNETs and their vasculature (Sodir et al., 2011). Prior microarray analyses of RABL6A depleted human pNET cell lines showed significant reduction in *c-Myc* mRNA levels (Hagen et al., 2014), suggesting that RABL6A may normally promote *c-Myc* transcription.

To determine if RABL6A promotes the expression of *c-Myc* mRNA in pancreatic islets, quantitative RT-PCR assays were performed on cDNA from the isolated islets of WT vs R6KO and RT2 vs RT2; R6KO mice. Significant downregulation of *c-Myc* expression was seen in normal islets from 8-10-week-old female and 12-week-old male R6KO mice lacking RABL6A compared to age-matched WT controls (***Figure 6A***). To assess the effect of RABL6A loss on *c-Myc* levels in neoplastic lesions, identical analyses were performed using transformed islets from RT2 and RT2; R6KO pancreata. The pancreatic perfusions from these animals yielded a mixture of normal, hyperplastic and angiogenic islets. As shown in ***Figure 6B*** and ***Table 1***, *c-Myc* mRNA expression was significantly reduced by ~2-fold in neoplastic islets from 8-10-week RT2; R6KO females vs RT2 controls (p<0.05). RABL6A deficiency in RT2; R6KO 12-week males likewise yielded a more than 3-fold average decrease (albeit not statistically significant,p=0.11) in *c-Myc* expression (***Figure 6B; Table 1***). These data suggest RABL6A promotes *c-Myc* mRNA expression in normal and transformed pancreatic islets.

**Figure 6.**
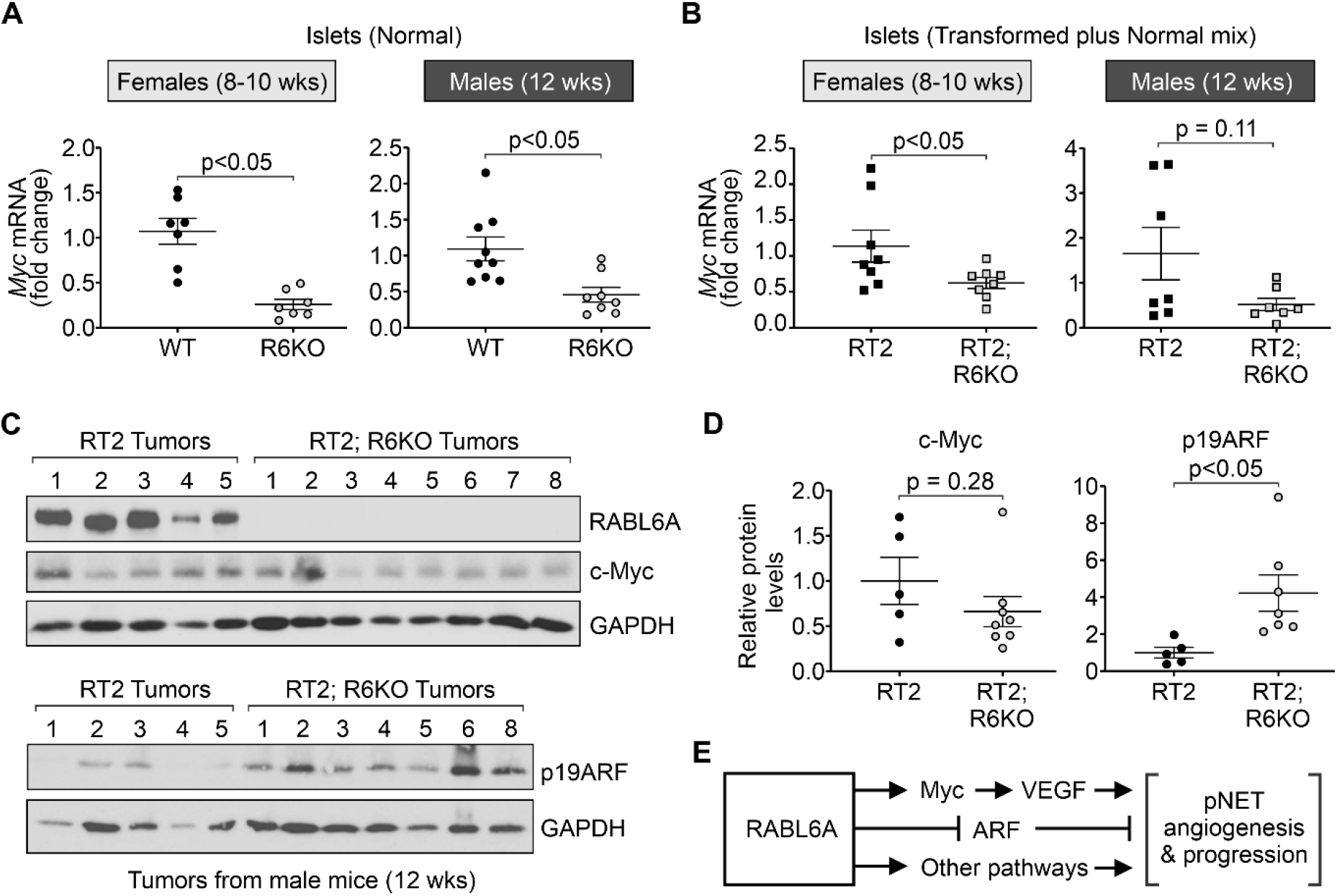
RABL6A promotes *c-Myc* expression in mouse pancreatic islets and tumors. **(A and B)** RNA was isolated from frozen islets (normal or a mixture of transformed plus normal), as indicated, and qRT-PCR was performed to compare the transcript levels of *c-Myc* between WT and R6KO **(A)** or RT2 and RT2; R6KO **(B)** cohorts. P values are shown. N = Females (WT = 7, RT2 = 8, R6KO = 7, RT2; R6KO = 8); Males (WT = 9, RT2 = 7, R6KO = 8, RT2; R6KO = 7). **(C)** Top: Western analysis of RABL6A, c-Myc, and GAPDH (loading control) in tumors from 12-week-old male RT2 and RT2; R6KO mice. Bottom: Western analysis of p19ARF and GAPDH in same tumor lysates (minus #7 since it was used up) run on separate gels. **(D)** ImageJ densitometric analysis of western data (from C) for relative levels of c-Myc and p19ARF in RT2; R6KO tumors relative to RT2 controls. N = c-Myc (RT2 = 5, RT2; R6KO = 8); p19ARF (RT2 = 5, RT2; R6KO = 7). **(E)** Schematic summarizing our findings and proposed working model for how RABL6A promotes pNET pathogenesis. Data in A, B and D are presented as mean +/- S.E.M. Linear regression was used to evaluate differences in transcript and protein levels between groups. Each dot represents an individual animal. **Source data 1:** Figure 6A source data. **Source data 2:** Figure 6B source data. **Source data 3:** Figure 6D source data.

Western analyses of isolated pNETs from RT2 versus RT2; R6KO mice were performed to determine if RABL6A deficiency caused reduced expression of c-Myc protein (***Figure 6C; Table 1***). Pancreatic tumors from 10-week female cohorts were either absent or too small to be examined by western blotting, consequently tumors from 12-week-old male mice were assayed. Immunoblotting for RABL6A verified its absence in RT2; R6KO tumors whereas high levels of expression were seen in RT2 lesions. Although results did not reach statistical significance, the majority of RT2; R6KO tumors expressed lower levels of c-Myc protein compared to RABL6A-positive RT2 tumors (***Figure 6C, D; Table 1***).

Prior studies in RT2 mice demonstrated an important role for the p19ARF tumor suppressor in reducing pNET development (Ulanet & Hanahan, 2010). Specifically, genetic deletion of *Arf* significantly accelerated tumor formation by promoting the angiogenic switch. Since RABL6A promotes pNET angiogenesis and physically associates with the ARF protein (Tompkins et al., 2006), we speculated RABL6A may impair ARF expression. The same tumor samples from ***Figure 6C*** (top panels) were examined by western blotting for mouse p19ARF (bottom panels, minus tumor #7 because sample was used up). As predicted, loss of RABL6A in RT2; R6KO tumors correlated with a significant, 4.2-fold average increase in p19ARF expression (***Figure 6C, D; Table 1***). These exciting findings support a model in which oncogenic RABL6A may promote pNET angiogenesis and progression through upregulation of c-Myc and/or downregulation of ARF in pancreatic islets (***Figure 6E***).

## Discussion

Molecular alterations driving pNET pathogenesis are only partly understood, but what is known has helped guide current patient therapies. Standard treatments such as everolimus (inhibitor of mammalian target of rapamycin [mTOR]) and sunitinib (inhibitor of receptor tyrosine kinases, including VEGFR) (Raymond et al., 2011; Strosberg et al., 2017; Yao et al., 2011) were supported by patient tumor profiling (Jiao et al., 2011; Missiaglia et al., 2010) and drug studies in RT2 mice demonstrating their effectiveness (Chiu et al., 2010; Olson et al., 2011). However, none of the current therapies for pNETs are curative and overall patient survival is not significantly improved. Greater insight into key drivers of the disease is still needed to inform new treatments with innovative targeted and/or combination therapies. This study explored the *in vivo* significance of a new player in pNET biology, RABL6A. We found that loss of RABL6A in the RT2 pNET model reduced tumor burden, angiogenesis and mitoses in an age-dependent manner that coincided with improved survival.

Several prior observations predicted that RABL6A would play an oncogenic role in pNET pathogenesis *in vivo*. First, RABL6A is upregulated at the genetic and protein level in patient pNETs (Hagen et al., 2014; Scott et al., 2020). Second, RABL6A promotes pNET cell survival and proliferation through multiple, clinically druggable pathways that include CDK4/6-RB1, PP2A and Akt/mTOR (Hagen et al., 2014; Umesalma et al., 2019). Third, microarray analyses of cultured pNET cells expressing or lacking RABL6A identified a RABL6A-regulated gene expression profile that included activation of c-Myc, VEGFR and EGFR pathways (Hagen et al., 2014), all of which are major mediators of tumor angiogenesis.

Angiogenesis is critical to pNET development and progression. This is evidenced by the clinical use of sunitinib to block pro-angiogenic receptor tyrosine kinases, like VEGFR, and their downstream effectors (PI3K/Akt/mTOR, PKCs, Ras/MAPK) in pNET therapy (Aparicio-Gallego et al., 2011; Raymond et al., 2011). Oncogenic c-Myc transcribes *VEGF* (Baudino et al., 2002), a VEGFR ligand, and is critical for pNET angiogenesis and progression (Casanovas et al., 2005; Inoue et al., 2002; Pelengaris et al., 2002; Sodir et al., 2011). We found downregulation of *c-Myc* transcripts in RABL6A-deficient pancreatic islets, in keeping with reduced angiogenic islets and vasculature in young RT2; R6KO mice. Those results support the conclusion that RABL6A may promote pNET angiogenesis in RT2 mice, at least in part, via c-Myc upregulation (as depicted in ***Figure 6E***).

Other RABL6A effectors likely contribute to its pro-angiogenic activity in pNET development. RABL6A promotes AKT-mTOR signaling via inactivation of the PP2A tumor suppressor (Umesalma et al., 2019), and numerous studies have shown that activated AKT-mTOR signaling promotes tumor angiogenesis (Karar & Maity, 2011). The role of PP2A in angiogenesis is less clear but a recent study of a vascular neoplasm, called hemangioma, revealed that inactivation of PP2A promotes angiogenesis and hemangioma formation (Xie et al., 2015). Notably, RABL6A physically interacts with the ARF tumor suppressor (Tompkins et al., 2006). Since ARF suppresses pNETs in RT2 mice by blocking the angiogenic switch (Ulanet & Hanahan, 2010), RABL6A may promote tumor angiogenesis by antagonizing ARF. That concurs with our finding that p19ARF protein expression is significantly reduced in RABL6A-expressing pNETs. It is possible that RABL6A, which is predominantly cytosolic but shuttles into the nucleus (Tompkins et al., 2006), mobilizes ARF from the nucleus and nucleoli (where it is most stable) into the cytosol where it undergoes proteasomal and lysosomal degradation (Colombo et al., 2005; den Besten, Kuo, Williams, & Sherr, 2005; Kuo, den Besten, Bertwistle, Roussel, & Sherr, 2004; Rodway, Llanos, Rowe, & Peters, 2004; Seo, Seong, Lee, Oh, & Song, 2020). Further investigations will be needed to test these possibilities.

The RT2 mouse model is broadly used in pNET research. It provides a reliable, fully penetrant model of multi-stage pNET progression that has been validated for preclinical drug testing. Besides demonstrating *in vivo* anti-tumor activity of rapamycin and sunitinib (Chiu et al., 2010; Olson et al., 2011), RT2 mice have helped rule out pathways with low clinical relevance in pNETs, such as IGF-1R (Reidy-Lagunes et al., 2012; Ulanet, Ludwig, Kahn, & Hanahan, 2010). Nevertheless, there are some limitations of this model. Inhibition of central tumor suppressors, p53 and RB1, by SV-40 large T-antigen in RT2 mice causes aggressive, highly proliferative pNETs. In this regard, the RT2 molecular signatures and phenotype more closely recapitulate high grade 3 (G3) insulinomas, which are well-differentiated pNETs that have a high proliferative index (Ki-67>20%). Moreover, the absence of functional RB1 likely reduced the extent to which RABL6A loss suppressed pNET formation since RABL6A functions partly through RB1 inhibition (Hagen et al., 2014). On the other hand, many low G1/G2 pNETs lack functional RB1 due to overexpression of CDK4 and CDK6 or loss of CDK inhibitors, like p16INK4a (Muscarella et al., 1998; Serrano et al., 2000; L. H. Tang et al., 2012). Thus, the RT2 model enabled us to determine the contribution of RABL6A to pNET development in a clinically relevant, RB1 inactive setting common in patient tumors. Notably, earlier drug studies in RT2 mice targeting c-Myc (Sodir et al., 2011), VEGF (Bergers et al., 2003; Casanovas et al., 2005), and PI3K/Akt/mTOR signaling (Soler et al., 2016), yielded similar phenotypes of reduced tumor progression and angiogenesis as seen in RT2; R6KO animals.

It will be important to explore the in vivo role of RABL6A in pNET pathogenesis using other relevant models of the disease. Several groups have modeled deficiency of the multiple endocrine neoplasia 1 (*Men1*) gene in mice using the rat insulin promoter to achieve inactivation in β cells (Bertolino et al., 2003; Crabtree et al., 2003). Patients with the MEN1 familial tumor syndrome develop multiple endocrine tumors, such as insulinomas, and somatic mutations of human *MEN1* are also common in various sporadic NETs, including those of the pancreas (Jensen, Berna, Bingham, & Norton, 2008; Jiao et al., 2011; Scarpa et al., 2017). In mice, loss of *Men1* alone yields insulinomas with high penetrance but long latency period (80-100% incidence by ~ 1 year of age) (Bertolino et al., 2003; Crabtree et al., 2003). The tumors develop in a multi-stage fashion similar to RT2 mice, with early formation of atypical hyperplastic islets progressing into angiogenic islets and ultimately low-grade neoplastic lesions, with no evidence of metastasis. Others recently demonstrated greatly accelerated formation of low-grade (G1/G2) insulinomas in mice with targeted inactivation of both *Men1* and *Pten*, with an onset of pNETs at 7 weeks of age (Wong et al., 2019). If RABL6A promotes tumor angiogenesis and development in these contexts, as expected, its loss would diminish the tumor phenotype. Because the *Rabl6* deficient mice used herein represent a global gene trap model, we cannot rule out potential contributions of small, N-terminal truncation products generated from exons 1-3 of the 15-exon *Rabl6* gene. At present, antibodies that recognize N-terminal sequences of RABL6A are lacking. If such mutant proteins are formed, they could diminish the *Rabl6* knockout phenotype if they retain some functional activity. Given these issues, conditional deletion of *Rabl6* in β cells is warranted to bypass limitations of the gene-trap system and tease apart specific contributions of RABL6A in pancreatic β cells versus the tumor microenvironment.

In general, more pronounced anti-tumoral effects of RABL6A loss were seen in female RT2 mice, potentially suggesting sex-dependent activities of RABL6A. However, the increase in survival associated with RABL6A deficiency was more apparent in RT2 males, arguing against a sex-dependent effect. It seems more likely that the timing of our analyses, chosen because of the different survival rates for female versus male RT2 mice, impacted our ability to detect effects of RABL6A loss on the tumor phenotype. In that regard, analyses in males were further spread out (8, 12, 16 weeks) than in females (8, 10, 12 weeks), lacking the 10-week time point where the most significant changes were observed in RT2; R6KO females.

The anti-tumor effects of RABL6A loss in RT2 mice were age dependent. The most significant reductions in tumor burden, angiogenesis or islet cell proliferation were observed in the earlier stages of pNET development, namely 8- to 10-week-old female or 12-week-old male RT2; R6KO mice. Conversely, the tumor suppressive effect of RABL6A-deficiency was lost in older animals as tumors progressed to advanced stages. Somewhat unexpectedly, 16-week-old RT2; R6KO males displayed greater mitotic content than RT2 counterparts and this coincided with a trend toward increased tumor burden. This may reflect development of resistance to RABL6A loss, enabling tumors to progress more aggressively. This mirrors an unwanted problem (drug resistance) that arises too often in the clinic during anti-cancer therapy, and it has been previously observed in RT2 drug studies. For example, treatment of RT2 mice with anti-angiogenic drugs, sunitinib or VEGFR2 inhibitor, causes temporary tumor regression followed by evasive drug resistance marked by increased invasiveness and malignancy. The resistance against anti-angiogenic therapy was mediated by upregulation of pro-angiogenic factors, FGF2 (Casanovas et al., 2005) and c-Met (Sennino, Ishiguro-Oonuma, Schriver, Christensen, & McDonald, 2013; Sennino et al., 2012). Similar mechanisms may have enabled RT2; R6KO tumors to circumvent the reduced c-Myc and upregulation of p19ARF caused by RABL6A loss, possibly facilitating the progression of tumors in older male mice.

In sum, this work demonstrates a significant role for RABL6A in driving pNET angiogenesis and development *in vivo*. RABL6A oncogenic activity was coincident with increased expression of *c-Myc* and reduced ARF in pancreatic islets and tumors. Together, these results highlight the potential benefit of pharmacologically targeting RABL6A pathways in pNET therapy and warrant further investigation into mechanisms of RABL6A action in pNET pathogenesis.

## Materials and Methods

### Mice

This mouse study tested the hypothesis that RABL6A loss by gene disruption will reduce pNET development in an established genetic model of the disease, RIP-Tag2 (abbreviated RT2). Animals were housed in barrier conditions. All mouse handling was conducted in strict compliance with The University of Iowa Institutional Animal Care and Use Committee (IACUC) policies (animal care protocol # 8111590), which adheres to the requirements of the National Institutes of Health Guide for the Care and Use of Laboratory Animals and the Public Health Service Policy on the Humane Care and Use of Laboratory Animals. All efforts were made to minimize animal suffering. Two RT2 male mice were kindly provided by Dr. Chris Harris (University of Rochester). Male RT2 mice (C57BL/6N strain) were bred with wild-type C57BL/6N females (Charles River) to obtain RT2 and WT experimental cohorts. RT2 males were mated with R6KO (i.e., *Rabl6^m/m^*, also on the C57BL/6N background) females to obtai
RT2; *R6R^+/m^* mice (50% incidence), which were further crossed with R6K0 mice to obtain RT2; R6KO (i.e., RT2; *Rabl6^m/m^*) mice (25% incidence). Breeding pairs of RT2; R6KO males and R6KO females were used to generate R6KO and RT2; R6KO experimental cohorts at 50:50 ratio. Mice were grouped by genotype and sex, and then randomly binned into the different experimental age groups for analyses. Animals that died in pen (DIP) or those which were euthanized for having low BCS conditions were used in survival analysis.

A sample size estimation was conducted when the study was designed. Numbers were based on power calculations, the timing and penetrance of the tumor phenotype in prior studies of the RT2 model (100% tumor penetrance by 14 weeks of age), logical scientific design and the prediction that RABL6A loss (R6KO) would delay the rate of tumor formation and extend survival. A cohort size of 30 animals per RT2 and RT2; R6KO genotype, yielding 10 mice for each of 3 time points, was projected using an F test for time effect on tumor size at a 5% significance level. Analyses were based on assumptions about anticipated tumor sizes from published data and the expectation that 10 mice per time point would represent 5 males and 5 females for each genotype. The final numbers differed from original estimates, ranging from 5-14 animals of each sex per genotype and time point for the different assays as well as expanded numbers for survival analyses (which ranged from 23 to 45 mice per sex and genotype). The change in sample sizes reflected 1) the observed >2-week difference in median survival between male and female RT2 mice, prompting different time points for the tumor analyses in female (8, 10 and 12 weeks) and male (8, 12 and 16 weeks) mice; 2) basis of the estimates on two factors (tumor burden and survival), but during the study other assays of tumor size (% endocrine area), vasculature, mitotic index and protein expression required addition of separate mice; 3) the Covid-19 shutdown prevented continued studies for most of 2020, except for maintenance of the colony. The final sample sizes allowed detection of statistically significant differences in survival, tumor formation, angiogenesis, mitoses and gene/protein expression between RT2 and RT2; R6KO mice.

### Genotyping

All mice were genotyped by PCR. Genomic DNA were extracted from tail snips of the new weanlings using RED Extract-N-Amp^TM^ PCR kit. Primers for *RT2* (+/tg) genotyping were obtained from NCI mouse repository: a) Transgene: forward-^5^’GGACAAACCACAACTAGAATGCAG^3^’, reverse-^5^’CAGAGCAGAATTGTGGAGTGG^3^’, and b) WT: forward-^5^’CACCGGAGAATGGGAAGCCGAA^3^’, reverse-^5^’TCCACACAGATGGAGCGTCCAG^3^’. *RT2* PCR cycling conditions: a) 94 °C 3 min, b) 35 cycles of 94 °C 30 sec, 60 °C 30 sec, 72 °C 30 sec, and c) 72 °C 3 min. *RABL6* genotyping was performed using a common forward primer-^5^’CTACAGGACCTGTGGTTGTCT^3^’, and two reverse primers-^5^’CTGGCTCTCATGGAATCGTG^3^’ (WT *RABL6* specific) and ^5^’CCAACTGACCTTGGGCAAGAACAT^3^’ (gene-trap mutant-specific). *RABL6* PCR cycling conditions: a) 95 °C 10 min, b) 35 cycles of 95 °C 15 sec, 56.5 °C 45 sec, 60 °C 1 min. PCR products were run on 1.5% agarose gel containing ethidium bromide and imaged using UVP BioSpectrum^®^ 610 Imaging System.

### Mouse blood withdrawal and Insulin ELISA

Mice were starved for five hours prior to blood withdrawal. Their lateral saphenous vein was punctured using a 25G7/8 needle to collect 30-40 μL blood per mouse into a heparinized capillary tube. The blood was immediately mixed with 1 μL of 5% EDTA to prevent clotting and later centrifuged at 5000 rcf for 5 min at 4 °C. The supernatant (plasma) was carefully collected using a pipette and stored at −80 °C. To measure plasma insulin levels, frozen plasma was thawed on ice and 5 uL/sample loaded in duplicate wells onto a 96 well plate in Mouse Ultrasensitive Insulin ELISA kit from ALPCO^®^. Absorbance of the final products was measured at 450 nm using BioTek^®^ Synergy 4 plate reader. Data analysis was performed using GraphPad Software.

### Islet isolation

Mice were euthanized by cervical dislocation following isoflurane-induced anesthesia in a glass chamber. Pancreatic islets of Langerhans (including tumorigenic islets) were isolated using pancreatic perfusion technique (Kang et al., 2018). 2-3 mL of 0.8 mg/mL collagenase in HBSS solution was perfused into the pancreas through the cannulated common bile duct (clamped at its hepatic branch). Inflated pancreas was carefully removed and digested at 37°C for 10 min. Islets and tumors were then released by gentle agitation, washed in RPMI with 1% FBS and purified on Histopaque 1077 and 1119 gradients. Islet pellets and tumors were flash frozen in liquid nitrogen and stored at −80°C.

### RNA isolation and qRT-PCR analysis

RNA was prepared from isolated islets using Qiagen RNAeasy^®^ Plus Mini Kit. 40 μL of 1M dithiothreitol (DTT) was added per mL of lysis buffer to inhibit abundantly expressed pancreatic ribonucleases (Chen, Ling, & Gallie, 2004). cDNA was synthesized from 100-200 ng RNA using SuperScript^®^ III First Strand cDNA preparation kit (Invitrogen). Diluted cDNA was then used for qPCR (cycling conditions: denaturation at 95°C for 10 min followed by 40 cycles of 95°C for 15 sec and 60°C for 1 min) with gene-specific primers in the presence of iQ^TM^ SYBR^®^ Green Supermix reagent on a Bio-Rad CFX96^TM^ Real-Time System. All samples were run in triplicate and fold changes in each gene mRNA levels were calibrated to *Hprt* mRNA expression and computed using the 2^-ΔΔCt^ method.

### qRT-PCR primers

The National Center of Biotechnology Information (NCBI) Primer-BLAST online tool was used to design primers specific for the mouse genes of interest. a) *Hprt* (housekeeping gene used for normalization): forward-^5’^GCCCCAAAATGGTTAaGgtTG^3’^, reverse-^5^TGGCCTGTATCCAACACTTC^3’^; and b) *c-Myc*: forward-^5’^CCTGTACCTCGTCCGATTCC^3’^, reverse-^5’^TTCTTGCTCTTCTTCAGAGTCG^3’^.

### Analysis of tumor burden

Tumors (lesions ≥ 1mm) were excised from a fully perfused pancreas and their dimensions measured with a ruler. Individual tumor volumes were calculated using the formula: volume = width^2^ × length × 0.52 and summed up to calculate tumor burden per mouse as previously described (Ulanet & Hanahan, 2010). Tumors were flash frozen in liquid nitrogen for western blot analysis.

### Histopathological analyses

Isolated pancreas tissues were fixed in formalin (10% neutral buffered formalin), paraffin-embedded, and processed for hematoxylin and eosin (H&E) staining. Histopathological scoring of tissues was performed in masked manner and by following principles and approaches of reproducible scoring analysis (Meyerholz & Beck, 2018). Endocrine area, angiogenic islets, vasculature, and number of mitoses were analyzed on H&E-stained specimens using a brightfield microscope. To compute percentage endocrine area, the area covered by islets (normal and tumorigenic) in three randomly selected pancreatic regions viewed at low (20x) magnification was divided by the total area (% endocrine area= Islet area / total pancreatic area x 100). The percentage of angiogenic islets was calculated as the number of angiogenic islets divided by the total (non-angiogenic plus angiogenic) multiplied by 100. To calculate the percent vasculature (i.e., vessels larger than capillaries), individual angiogenic islets were observed at 100x magnification and the ratio of area covered by blood vessel to the total area of an islet multiplied by 100 (% vasculature = vessel area / islet area x 100). Number of mitoses were counted on 3-4 random fields of view within an islet at 600x magnification. Percentage vasculature and number of mitoses were quantified on 3-4 randomly selected enlarged islets in the pancreas of RT2 and RT2; R6KO mice.

### Western blotting and antibodies

Flash frozen tumors were stored at −80^°^C and pulverized in LN2 using a mortar and pestle. Samples were lysed for 30min on ice with RIPA buffer (50 mM Tris, pH 8.0, 150 mM NaCl, 1% Triton X-100, 0.1% SDS, 0.5% sodium deoxycholate) containing 1 mM NaF, protease and phosphatase inhibitor cocktails (Sigma, P-8340 & P-0044) as well as 30 μM phenylmethylsulfonyl fluoride (PMSF). Protein concentrations of the extracts were determined by BCA protein assay (ThermoFisher, Cat. # 23228). Equivalent amounts of protein per sample were separated by SDS-PAGE and transferred onto PVDF membranes (Millipore). Membranes were blocked with 5% non-fat milk or 5% BSA in TBST (Tris-buffered saline containing Tween-20) depending on the antibody. Proteins were detected using HRP-conjugated secondary antibodies and enhanced chemiluminescence (ECL, Amersham, Buckinghamshire, UK). Densitometry quantification was performed using ImageJ (NIH). Antibodies were used according to supplier guidelines and included: GAPDH mouse monoclonal (no. Ab8245) from Abcam; Myc [Y69] rabbit monoclonal (no. Ab32072) from Abcam; HRP-coupled secondary antibodies (nos. NA934 and NA935) from Sigma; and RABL6A polyclonal rabbit antibody produced in the Quelle laboratory (Muniz et al., 2013; Tompkins et al., 2006).

### Statistics

Western data were imaged by scanning densitometry and quantified by ImageJ (NIH). Quantified data were presented as the mean +/- SEM for the biological replicates in each analysis. Linear regression was utilized to evaluate differences in tumor volume, average number of number of mitoses, transcript levels and protein levels; a log-transformation was applied to each outcome prior to analysis to meet model assumptions. Beta regression models were utilized to evaluated differences in the proportion of endocrine area, angiogenic islets, and vascular area per islet. The Kaplan-Meier method was used to estimate the survival curves, and group comparisons were made using the log rank test. Cox regression was used to evaluate the overall effect of genotype while adjusting for sex. Linear mixed effects regression models were used to estimate and compare group-specific changes in plasma insulin over time; a log-transformation was applied to stabilize the variance. Accounting for unequal variances or random variability attributable to repeat measurements within a mouse was done, as necessary. All tests were two-sided and assessed at the 5% significance level using SAS v9.4 (SAS Institute, Cary, NC).

## Author Contributions

Study conceptualization and design: CKM, BWD, SS, DKM, and DEQ. Experimentation, data acquisition and analysis: CKM, CAK, VPM, CB, SM, KDZ, PB and MRL. Supervision and scientific interpretation of data: BWD, SS, DKM, DEQ. All authors reviewed and approved the manuscript.

## Acknowledgements

We thank Drs. Fred Quelle, Rory Fisher, Adam Dupuy, and Chandrikha Chandrasekharan for their advice and ideas contributing the advancement of this project. We also thank Dr. Chris Harris for providing RT2 breeders, Jussara Hagen for technical assistance generating R6KO mice, and the Iowa NET SPORE group for their critical comments and feedback. This project was supported by an NCI Neuroendocrine Tumor SPORE P50 CA174521 [Project 2, DEQ and BWD], an NCI Core Grant P30 CA086862 [University of Iowa Holden Comprehensive Cancer Center], and several University of Iowa Graduate College fellowships [CKM].

## Supplementary Figures

**Figure 1- figure supplement 1.**
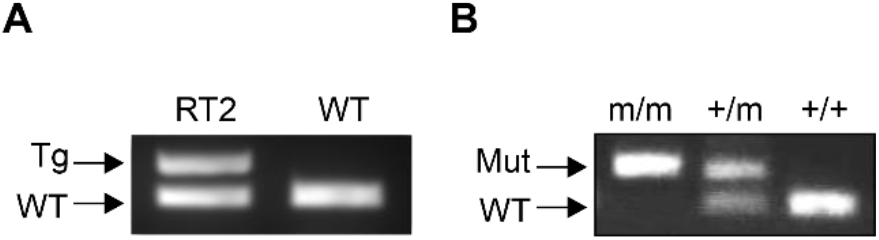
Genotyping RIP-Tag2 (RT2) and R6KO mice. **(A)** PCR gel electrophoresis image shows bands representing SV40 large T-antigen (Tg) or a WT DNA segment in RT2 and WT mice. **(B)** Gel images showing bands for WT or mutant *Rabl6* alleles in R6KO (m/m), het (+/m) and WT (+/+) mouse genomic DNA.

**Figure 3- figure supplement 1.**
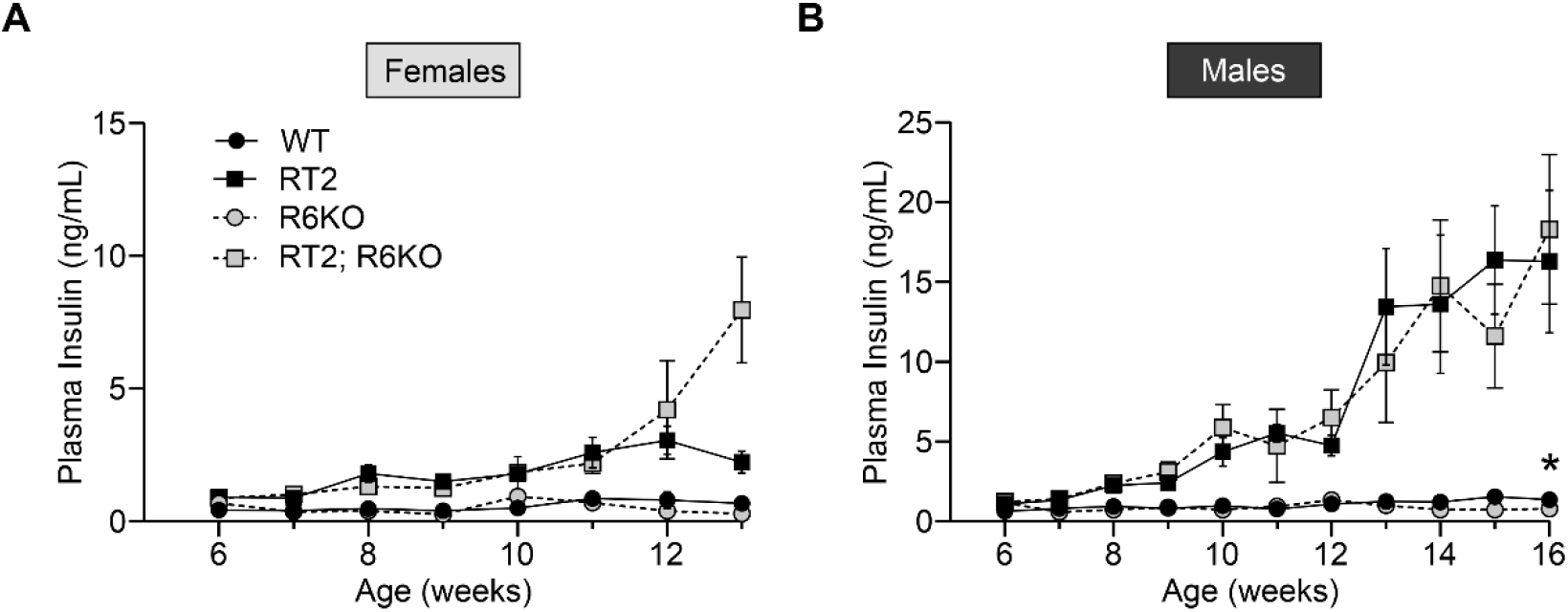
Graphs depicting plasma insulin concentration in female (6-13 weeks) and male (6-16 weeks) experimental mice. Plasma insulin was measured by using Mouse Ultrasensitive Insulin ELISA kit from ALPCO^®^. (A and B) Plasma insulin in WT and R6KO animals is low. WT males have significantly higher plasma insulin concentration than R6KO males (*p<0.05), however, no difference was observed between WT and R6KO females (p = 0.10). Both RT2 and RT2; R6kO females (A) and males (B) exhibit gradual increase in plasma insulin levels as they grow older with no observable difference between the genotypes (p = 0.19 and 0.58 respectively). Data are presented as mean +/- SEM. Linear mixed effects regression models were used for statistical comparison of insulin concentration between groups. **Source data 1:** Figure 3- figure supplement 1 source data

**Figure 3- figure supplement 2.**
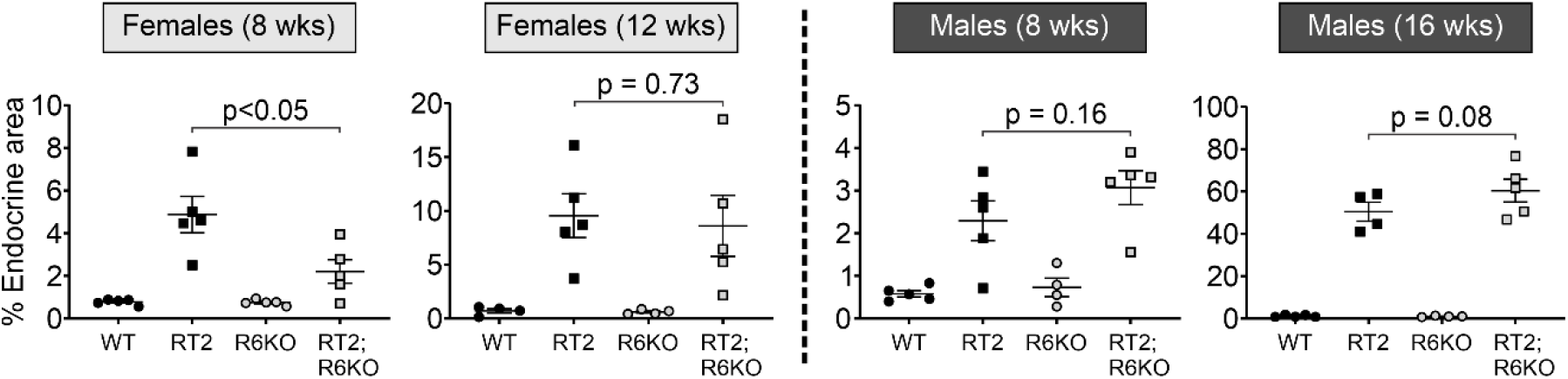
Percent endocrine area in the pancreata of female (left) and male (right) experimental mice at the indicated ages. 8-week-old RT2; R6KO females have significantly reduced endocrine area compared to the RT2 controls (p<0.05), whereas no difference is seen between the groups at 12 weeks of age (p = 0.73). Pancreata of RT2 and RT2; R6KO males at 8 and 16 weeks do not show differences in percent endocrine area (p = 0.16 and p=0.08, respectively). Data are presented as mean +/- SEM with each dot representing an individual mouse. N = Females: 8 weeks (WT = 5, RT2 = 5, R6KO = 4, RT2; R6KO = 5), 12 weeks (WT = 4, RT2 = 5, R6KO = 4, RT2; R6KO = 5); Males: 8 weeks (WT = 5, RT2 = 5, R6KO = 4, RT2; R6KO = 5), 16 weeks (WT = 5, RT2 = 4, R6KO = 4, RT2; R6KO = 5). Beta regression models were used for statistical analysis. **Source data 1:** Figure 3- figure supplement 2 source data

**Figure 4- figure supplement 1.**
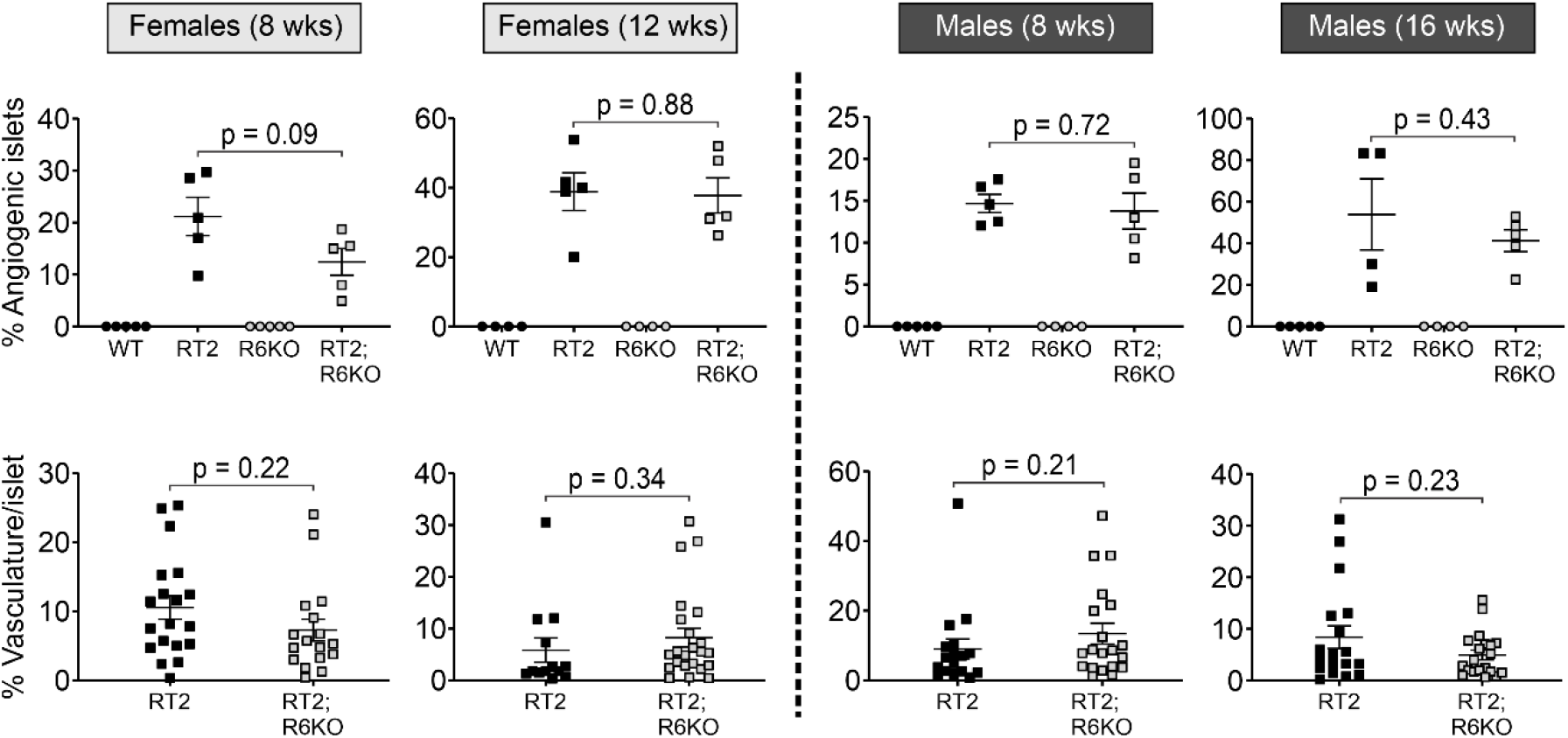
Scatterplots showing the percentage of angiogenic islets per pancreas (top row) or percent vascular area per angiogenic islet (bottom row) in female (left) and male (right) mice of the indicated genotypes. Data represent the mean +/- SEM. Each dot represents an individual mouse for endocrine area analyses, whereas an individual islet for vascular area analyses. N = Females: 8 weeks (WT = 5, RT2 = 5, R6KO = 4, RT2; R6KO = 5), 12 weeks (WT = 4, RT2 = 5, R6KO = 4, RT2; R6KO = 5); Males: 8 weeks (WT = 5, RT2 = 5, R6KO = 4, RT2; R6KO = 5), 16 weeks (WT = 5, RT2 = 4, R6KO = 4, RT2; R6KO = 5). Beta regression models were used for statistical analysis. **Source data 1:** Figure 4- figure supplement 1 source data

**Figure 5- figure supplement 1.**
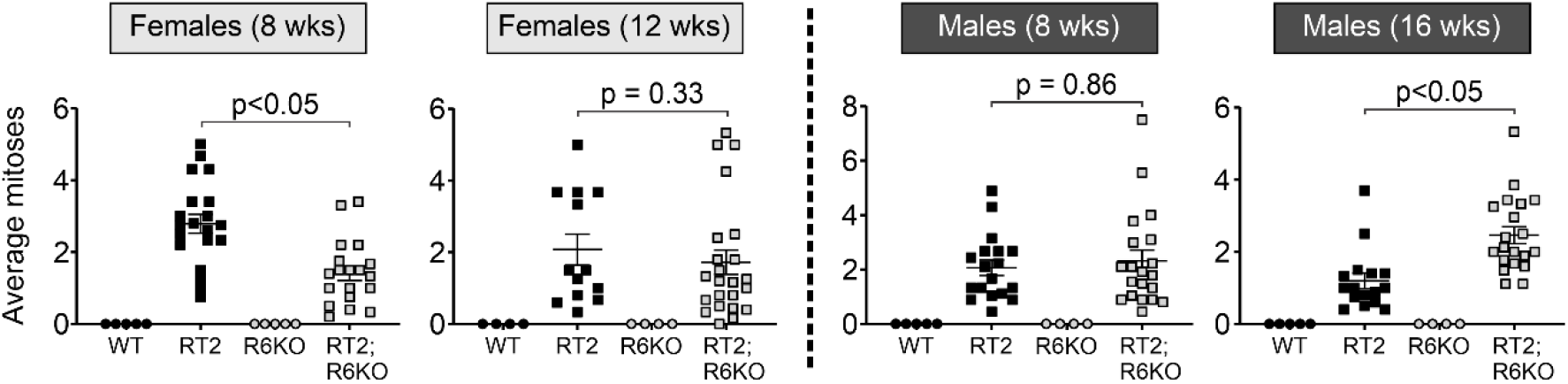
Scatterplots depicting the average number of mitoses per field of vision at 600x magnification for individual female (left) and male (right) pancreatic islets. Transformed islets of RT2; R6KO females (8 weeks) have significantly reduced number of mitoses compared to those of RT2 females (p<0.05). 16-week-old RT2; R6KO males have higher number of mitoses compared to their RT2 counterparts (p<0.05). Data represent the mean +/- SEM. Each dot represents an individual islet: normal islets in WT and R6KO mice versus transformed (hyperplastic, angiogenic, or tumor) islets in RT2 and RT2; R6KO mice. N = Females: 8 weeks (WT = 5, RT2 = 5, R6KO = 4, RT2; R6KO = 5), 12 weeks (WT = 4, RT2 = 5, R6KO = 4, RT2; R6KO = 5); Males: 8 weeks (WT = 5, RT2 = 5, R6KO = 4, RT2; R6KO = 5), 16 weeks (WT = 5, RT2 = 4, R6KO = 4, RT2; R6KO = 5). Linear regression was used to evaluate differences in the average number of mitoses. **Source data 1:** Figure 5- figure supplement 1 source data

## References

Aparicio-Gallego, G., Blanco, M., Figueroa, A., García-Campelo, R., Valladares-Ayerbes, M., Grande-Pulido, E., & Antón-Aparicio, L. (2011). New insights into molecular mechanisms of sunitinib-associated side effects. Mol Cancer Ther, 10(12), 2215–2223.

Baudino, T. A., McKay, C., Pendeville-Samain, H., Nilsson, J. A., Maclean, K. H., White, E. L.,… Cleveland, J. L. (2002). c-Myc is essential for vasculogenesis and angiogenesis during development and tumor progression. Genes & Development, 16(19), 2530–2543.

Bergers, G., Song, S., Meyer-Morse, N., Bergsland, E., & Hanahan, D. (2003). Benefits of targeting both pericytes and endothelial cells in the tumor vasculature with kinase inhibitors. J Clin Invest, 111(9), 1287–1295.

Bertolino, P., Tong, W. M., Herrera, P. L., Casse, H., Zhang, C. X., & Wang, Z. Q. (2003). Pancreatic beta-cell-specific ablation of the multiple endocrine neoplasia type 1 (MEN1) gene causes full penetrance of insulinoma development in mice. Cancer Res, 63(16), 4836–4841.

Casanovas, O., Hicklin, D. J., Bergers, G., & Hanahan, D. (2005). Drug resistance by evasion of antiangiogenic targeting of VEGF signaling in late-stage pancreatic islet tumors. Cancer Cell, 8(4), 299–309.

Chen, Z., Ling, J., & Gallie, D. R. (2004). RNase activity requires formation of disulfide bonds and is regulated by the redox state. Plant Mol Biol, 55(1), 83–96.

Chiu, C. W., Nozawa, H., & Hanahan, D. (2010). Survival Benefit With Proapoptotic Molecular and Pathologic Responses From Dual Targeting of Mammalian Target of Rapamycin and Epidermal Growth Factor Receptor in a Preclinical Model of Pancreatic Neuroendocrine Carcinogenesis. Journal of Clinical Oncology, 28(29), 4425–4433.

Christofori, G., Naik, P., & Hanahan, D. (1994). A second signal supplied by insulin-like growth factor II in oncogene-induced tumorigenesis. Nature, 369(6479), 414–418.

Colombo, E., Bonetti, P., Lazzerini Denchi, E., Martinelli, P., Zamponi, R., Marine, J. C.,… Pelicci, P. G. (2005). Nucleophosmin is required for DNA integrity and p19Arf protein stability. Mol Cell Biol, 25(20), 8874–8886.

Crabtree, J. S., Scacheri, P. C., Ward, J. M., McNally, S. R., Swain, G. P., Montagna, C.,… Collins, F. S. (2003). Of mice and MEN1: Insulinomas in a conditional mouse knockout. Mol Cell Biol, 23(17), 6075–6085.

Dasari, A., Shen, C., Halperin, D., Zhao, B., Zhou, S., Xu, Y.,… Yao, J. C. (2017). Trends in the Incidence, Prevalence, and Survival Outcomes in Patients With Neuroendocrine Tumors in the United States. Jama Oncology, 3(10), 1335–1342.

den Besten, W., Kuo, M. L., Williams, R. T., & Sherr, C. J. (2005). Myeloid leukemia-associated nucleophosmin mutants perturb p53-dependent and independent activities of the Arf tumor suppressor protein. Cell Cycle, 4(11), 1593–1598.

Feng, Y., Yan, S., Huang, Y., Huang, Q., Wang, F., & Lei, Y. (2020). High expression of RABL6 promotes cell proliferation and predicts poor prognosis in esophageal squamous cell carcinoma. BMC Cancer, 20(1), 602.

Folkman, J., Watson, K., Ingber, D., & Hanahan, D. (1989). Induction of angiogenesis during the transition from hyperplasia to neoplasia. Nature, 339(6219), 58–61.

Hagen, J., Muniz, V. P., Falls, K. C., Reed, S. M., Taghiyev, A. F., Quelle, F. W.,… Quelle, D. E. (2014). RABL6A promotes G1-S phase progression and pancreatic neuroendocrine tumor cell proliferation in an Rb1-dependent manner. Cancer Res, 74(22), 6661–6670.

Halfdanarson, T. R., Rabe, K. G., Rubin, J., & Petersen, G. M. (2008). Pancreatic neuroendocrine tumors (PNETs): incidence, prognosis and recent trend toward improved survival. Annals of Oncology, 19(10), 1727–1733.

Hanahan, D. (1985). HERITABLE FORMATION OF PANCREATIC BETA-CELL TUMORS IN TRANSGENIC MICE EXPRESSING RECOMBINANT INSULIN SIMIAN VIRUS-40 ONCOGENES. Nature, 315(6015), 115–122.

Hanahan, D., & Folkman, J. (1996). Patterns and emerging mechanisms of the angiogenic switch during tumorigenesis. Cell, 86(3), 353–364.

Inoue, M., Hager, J. H., Ferrara, N., Gerber, H. P., & Hanahan, D. (2002). VEGF-A has a critical, nonredundant role in angiogenic switching and pancreatic beta cell carcinogenesis. Cancer Cell, 1(2), 193–202.

Jensen, R. T., Berna, M. J., Bingham, D. B., & Norton, J. A. (2008). Inherited pancreatic endocrine tumor syndromes: advances in molecular pathogenesis, diagnosis, management, and controversies. Cancer, 113(7 Suppl), 1807–1843.

Jiao, Y., Shi, C., Edil, B. H., de Wilde, R. F., Klimstra, D. S., Maitra, A.,… Papadopoulos, N. (2011). DAXX/ATRX, MEN1, and mTOR Pathway Genes Are Frequently Altered in Pancreatic Neuroendocrine Tumors. Science, 331(6021), 1199–1203.

Kang, C., Xie, L., Gunasekar, S. K., Mishra, A., Zhang, Y., Pai, S.,… Sah, R. (2018). SWELL1 is a glucose sensor regulating β-cell excitability and systemic glycaemia. Nature communications, 9(1), 367–367.

Karar, J., & Maity, A. (2011). PI3K/AKT/mTOR Pathway in Angiogenesis. Front Mol Neurosci, 4, 51.

Kohlmeyer, J. L., Kaemmer, C. A., Pulliam, C., Maharjan, C. K., Samayoa, A. M., Major, H. J.,… Quelle, D. E. (2020). RABL6A Is an Essential Driver of MPNSTs that Negatively Regulates the RB1 Pathway and Sensitizes Tumor Cells to CDK4/6 Inhibitors. Clinical Cancer Research, 26(12), 2997–3011.

Kuo, M. L., den Besten, W., Bertwistle, D., Roussel, M. F., & Sherr, C. J. (2004). N-terminal polyubiquitination and degradation of the Arf tumor suppressor. Genes & Development, 18(15), 1862–1874.

Li, Y.-Y., Fu, S., Wang, X.-P., Wang, H.-Y., Zeng, M.-S., & Shao, J.-Y. (2013). Down-Regulation of C9orf86 in Human Breast Cancer Cells Inhibits Cell Proliferation, Invasion and Tumor Growth and Correlates with Survival of Breast Cancer Patients. Plos One, 8(8), e71764.

Lui, K., An, J., Montalbano, J., Shi, J., Corcoran, C., He, Q.,… Huang, Y. (2013). Negative regulation of p53 by Ras superfamily protein RBEL1A. Journal of Cell Science, 126(11), 2436–2445.

Meyerholz, D. K., & Beck, A. P. (2018). Principles and approaches for reproducible scoring of tissue stains in research. Lab Invest, 98(7), 844–855.

Missiaglia, E., Dalai, I., Barbi, S., Beghelli, S., Falconi, M., della Peruta, M.,… Scarpa, A. (2010). Pancreatic Endocrine Tumors: Expression Profiling Evidences a Role for AKT-mTOR Pathway. Journal of Clinical Oncology, 28(2), 245–255.

Montalbano, J., Lui, K., Sheikh, M. S., & Huang, Y. (2009). Identification and Characterization of RBEL1 Subfamily of GTPases in the Ras Superfamily Involved in Cell Growth Regulation. Journal of Biological Chemistry, 284(27), 18129–18142.

Muniz, V. P., Askeland, R. W., Zhang, X., Reed, S. M., Tompkins, V. S., Hagen, J.,… Quelle, D. E. (2013). RABL6A Promotes Oxaliplatin Resistance in Tumor Cells and Is a New Marker of Survival for Resected Pancreatic Ductal Adenocarcinoma Patients. Genes & Cancer, 4(7-8), 273–284.

Muscarella, P., Melvin, W. S., Fisher, W. E., Foor, J., Ellison, E. C., Herman, J. G.,… Weghorst, C. M. (1998). Genetic alterations in gastrinomas and nonfunctioning pancreatic neuroendocrine tumors: an analysis of p16/MTS1 tumor suppressor gene inactivation. Cancer Res, 58(2), 237–240.

Olson, P., Chu, G. C., Perry, S. R., Nolan-Stevaux, O., & Hanahan, D. (2011). Imaging guided trials of the angiogenesis inhibitor sunitinib in mouse models predict efficacy in pancreatic neuroendocrine but not ductal carcinoma. Proceedings of the National Academy of Sciences, 108(49), E1275–1284.

Parangi, S., Dietrich, W., Christofori, G., Lander, E. S., & Hanahan, D. (1995). Tumor Suppressor Loci on Mouse Chromosomes 9 and 16 Are Lost at Distinct Stages of Tumorigenesis in a Transgenic Model of Islet Cell Carcinoma. Cancer Research, 55(24), 6071–6076.

Pelengaris, S., Khan, M., & Evan, G. I. (2002). Suppression of Myc-Induced Apoptosis in β Cells Exposes Multiple Oncogenic Properties of Myc and Triggers Carcinogenic Progression. Cell, 109(3), 321–334.

Raymond, E., Dahan, L., Raoul, J.-L., Bang, Y.-J., Borbath, I., Lombard-Bohas, C.,… Ruszniewski, P. (2011). Sunitinib Malate for the Treatment of Pancreatic Neuroendocrine Tumors. New England Journal of Medicine, 364(6), 501–513.

Reidy-Lagunes, D. L., Vakiani, E., Segal, M. F., Hollywood, E. M., Tang, L. H., Solit, D. B.,… Saltz, L. B. (2012). A phase 2 study of the insulin-like growth factor-1 receptor inhibitor MK-0646 in patients with metastatic, well-differentiated neuroendocrine tumors. Cancer, 118(19), 4795–4795.

Rindi, G., Klimstra, D. S., Abedi-Ardekani, B., Asa, S. L., Bosman, F. T., Brambilla, E.,… Cree, I. A. (2018). A common classification framework for neuroendocrine neoplasms: an International Agency for Research on Cancer (IARC) and World Health Organization (WHO) expert consensus proposal. Mod Pathol, 31(12), 1770–1786.

Rodway, H., Llanos, S., Rowe, J., & Peters, G. (2004). Stability of nucleolar versus non-nucleolar forms of human p14ARF. Oncogene, 23(37), 6186–6192.

Scarpa, A., Chang, D. K., Nones, K., Corbo, V., Patch, A. M., Bailey, P.,… Grimmond, S. M. (2017). Whole-genome landscape of pancreatic neuroendocrine tumours. Nature, 543(7643), 65–71.

Scott, A. T., & Howe, J. R. (2019). Evaluation and Management of Neuroendocrine Tumors of the Pancreas. Surg Clin North Am, 99(4), 793–814.

Scott, A. T., Weitz, M., Breheny, P. J., Ear, P. H., Darbro, B., Brown, B. J.,… Howe, J. R. (2020). Gene Expression Signatures Identify Novel Therapeutics for Metastatic Pancreatic Neuroendocrine Tumors. Clin Cancer Res.

Sennino, B., Ishiguro-Oonuma, T., Schriver, B. J., Christensen, J. G., & McDonald, D. M. (2013). Inhibition of c-Met reduces lymphatic metastasis in RIP-Tag2 transgenic mice. Cancer Res, 73(12), 3692–3703.

Sennino, B., Ishiguro-Oonuma, T., Wei, Y., Naylor, R. M., Williamson, C. W., Bhagwandin, V.,… McDonald, D. M. (2012). Suppression of tumor invasion and metastasis by concurrent inhibition of c-Met and VEGF signaling in pancreatic neuroendocrine tumors. Cancer Discov, 2(3), 270–287.

Seo, J., Seong, D., Lee, S. R., Oh, D. B., & Song, J. (2020). Post-Translational Regulation of ARF: Perspective in Cancer. Biomolecules, 10(8).

Serrano, J., Goebel, S. U., Peghini, P. L., Lubensky, I. A., Gibril, F., & Jensen, R. T. (2000). Alterations in the p16INK4a/CDKN2A tumor suppressor gene in gastrinomas. The Journal of clinical endocrinology and metabolism, 85(11), 4146–4156.

Sodir, N. M., Swigart, L. B., Karnezis, A. N., Hanahan, D., Evan, G. I., & Soucek, L. (2011). Endogenous Myc maintains the tumor microenvironment. Genes Dev, 25(9), 907–916.

Soler, A., Figueiredo, A. M., Castel, P., Martin, L., Monelli, E., Angulo-Urarte, A.,… Graupera, M. (2016). Therapeutic Benefit of Selective Inhibition of p110alpha PI3-Kinase in Pancreatic Neuroendocrine Tumors. Clin Cancer Res, 22(23), 5805–5817.

Strosberg, J. R., Halfdanarson, T. R., Bellizzi, A. M., Chan, J. A., Dillon, J. S., Heaney, A. P.,… Bergsland, E. K. (2017). The North American Neuroendocrine Tumor Society Consensus Guidelines for Surveillance and Medical Management of Midgut Neuroendocrine Tumors. Pancreas, 46(6), 707–714.

Sun, J. (2017). Pancreatic neuroendocrine tumors. Intractable & rare diseases research, 6(1), 21–21.

Tang, H., Ji, F., Sun, J., Xie, Y., Xu, Y., & Yue, H. (2016). RBEL1 is required for osteosarcoma cell proliferation via inhibiting retinoblastoma 1. Mol Med Rep, 13(2), 1275–1280.

Tang, L. H., Contractor, T., Clausen, R., Klimstra, D. S., Du, Y.-C. N., Allen, P. J.,… Harris, C. R. (2012). Attenuation of the Retinoblastoma Pathway in Pancreatic Neuroendocrine Tumors Due to Increased Cdk4/Cdk6. Clinical Cancer Research, 18(17), 4612–4620.

Tompkins, V., Hagen, J., Zediak, V. P., & Quelle, D. E. (2006). Identification of novel ARF binding proteins by two-hybrid screening. Cell Cycle, 5(6), 641–646.

Ulanet, D. B., & Hanahan, D. (2010). Loss of p19<sup>Arf</sup> Facilitates the Angiogenic Switch and Tumor Initiation in a Multi-Stage Cancer Model via p53-Dependent and Independent Mechanisms. Plos One, 5(8), e12454.

Ulanet, D. B., Ludwig, D. L., Kahn, C. R., & Hanahan, D. (2010). Insulin receptor functionally enhances multistage tumor progression and conveys intrinsic resistance to IGF-1R targeted therapy. Proceedings of the National Academy of Sciences, 107(24), 10791–10798.

Umesalma, S., Kaemmer, C. A., Kohlmeyer, J. L., Letney, B., Schab, A. M., Reilly, J. A.,… Quelle, D. E. (2019). RABL6A inhibits tumor-suppressive PP2A/AKT signaling to drive pancreatic neuroendocrine tumor growth. J Clin Invest, 130, 1641–1653.

Wong, C., Tang, L. H., Davidson, C., Vosburgh, E., Chen, W., Foran, D. J.,… Xu, E. Y. (2019). Two well-differentiated pancreatic neuroendocrine tumor mouse models. Cell Death & Differentiation.

Xie, F., Bao, X., Yu, J., Chen, W., Wang, L., Zhang, Z., & Xu, Q. (2015). Disruption and inactivation of the PP2A complex promotes the proliferation and angiogenesis of hemangioma endothelial cells through activating AKT and ERK. Oncotarget, 6(28), 25660–25676.

Yao, J. C., Shah, M. H., Ito, T., Bohas, C. L., Wolin, E. M., Van Cutsem, E… T, R. A. D. A. N. T. (2011). Everolimus for Advanced Pancreatic Neuroendocrine Tumors. New England Journal of Medicine, 364(6), 514–523.

